# Unified surface and volumetric projection of physiological imaging data

**DOI:** 10.1101/2022.01.28.477071

**Authors:** Thomas F. Kirk, Martin S. Craig, Michael A. Chappell

## Abstract

Projection of volumetric data onto the cortical surface is an important precursor to performing surface-based analysis. Numerous projection methods have been reported in the literature, many of which make assumptions which tie them to use with specific modalities, notably blood oxygenation level dependent (BOLD) imaging. This means that they may not be appropriate for use with modalities where subcortical tissue contributes a signal of interest. This work details a new projection that provides a number of generalisations and extensions to existing methods. Namely, it may be used to project arbitrary data without making modality-specific assumptions and can produce unified surface and volumetric representations of data (a concept also known as *grayordinates* space). When constrained to the same assumptions as existing methods, a comparison using simulation data shows that it produces similar outputs. When these assumptions are relaxed to project simulation data containing both cortical and subcortical signals to and from a unified surface and volume space, substantial and statistically significant differences in recovery of ground truth are observed compared to existing methods.

## 1. Introduction

The cerebral cortex has a thin and highly folded structure that may be thought of as a two-dimensional surface. Many properties of the cortex, notably the organisation of cortical areas, are best understood in the context of this surface topology and geometry [1]. In particular, on the assumption that the function of cortical areas will correlate closer according to geodesic (along the surface) than geometric (straight line) distance [2], surface-based analysis strategies should be inherently better suited to the study of the cortex than volumetric strategies, which divide the region of interest into voxels with no regard to underlying anatomy. Reported benefits of surface-based analysis of physiological imaging data include reduced bias and variance in kinetic modelling of positron emission tomography (PET) data [3]; improved localisation of functional areas within the cortex for blood oxygen level dependent (BOLD) magnetic resonance imaging (MRI) data [4, 5]; and improved inter-subject registration for multi-modal imaging [5, 6].

All surface-based analysis of volumetric data must start with a means of forward projecting data from the volume ‘onto the surface’. This is a complex operation for which there is no single analytic solution; instead, the methods that exist encode various assumptions about anatomy and the signal generation mechanism for the modality in question. Particularly important is the existence of the partial volume effect (PVE), a consequence of the low spatial resolution of imaging voxels in relation to the thickness of the cortex, which results in multiple tissues being present in a given voxel [7, 8]. PVE in voxels intersecting the cortex means that the acquired signal is likely to be a mixture of signal from cortical grey matter (GM), subcortical white matter (WM) and cerebrospinal fluid (CSF). Hence, a desirable feature of any projection method is that it is partial volume aware and attempts to minimise the amount of non-cortical signal that is projected onto the surface.

The majority of existing projection methods have been developed for use with BOLD data, which has important implications for how they treat WM signal. For example, Grova *et al*. expressly ignored WM signal in their method [9], whereas Operto *et al*. started by deriving the expected distribution of the BOLD signal on either side of the cortical ribbon, including in subcortical WM [10]. Similarly, the method developed by Glasser *et al*. is intended to be followed by a regression to remove any WM signal that *has* been projected to the surface [11]. These characteristics mean that these methods may not be suitable for the projection of data that does contain a WM signal of interest, and/or a WM signal that is spatially varying, for example that produced by arterial spin labelling (ASL).

Due to the fact that a small number of voxels in a typical volumetric dataset intersect the cortex, a surface-based analysis strategy will by design ignore the vast majority of imaging voxels. This is of no consequence if the cortex is the sole anatomy of interest, but if this is not the case, then a separate analysis must be run on subcortical voxels (likely a volumetric analysis, because the subcortex is not surface-like). Such a dual surface and volumetric analysis strategy has, for example, been adopted by the Human Connectome Project (HCP) [12, 11]: from a single timeseries of BOLD data, a conventional volumetric analysis is performed for the subcortex, whilst a surface analysis is performed for the cortex on a surface-projected copy of the data, maximising the extraction of useful information. The results of these separate analyses are then unified in *grayordinates* space, a domain of representation that allows data to be recorded both on the surface for the cortex and in the volume for the subcortex^1^. A notable aspect of this strategy is the bifurcation that is required: whilst the point of departure is a single volumetric timeseries, and the end-point is a single set of analysis results in grayordinates space, *two* parallel processing pipelines are required in between.

The objective of this work was to build a projection framework, which has been incorporated into the exiting Toblerone software package^2^, that extends existing methods in the following manner. Firstly, by providing the ability to perform forward (volume to surface) and reverse (surface to volume) projection, something that is not true of all existing methods. Secondly, by permitting the projection of arbitrary data, without making modality-specific assumptions, in particular data containing WM signal. Finally, by expanding the scope of projection to include unified surface and volumetric data representations, *i*.*e*., projection to and from a grayordinates-like space.

## 2. Theory

### 2.1. Hybrid space

Hybrid space is a generalisation of the HCP’s concept of grayordinates. It is a domain of data representation formed by taking the union of the surface and volume spaces, such that data in hybrid space has both a vertex-wise (surface) and voxel-wise (volume) representation for the cortex and subcortex respectively. For any space, be it surface, volumetric or hybrid, the spatial locations at which data values are recorded are hereafter referred to as *nodes*. Thus, in hybrid space, the set of nodes on which signal values are recorded comprises all surface vertices and all voxels of interest, as illustrated in figure 1.

**Figure 1:**
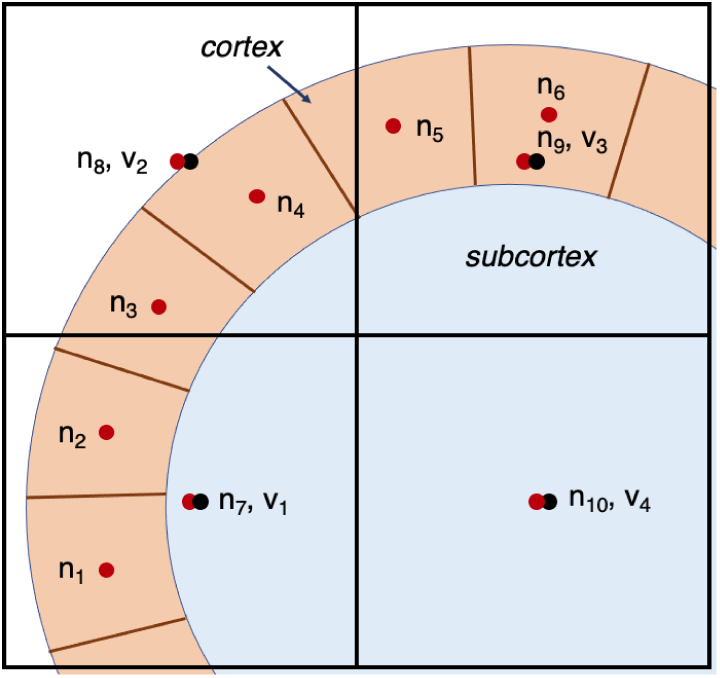
Schematic representation of voxels and nodes in hybrid space. The primitives comprising the cortical ribbon are denoted with brown boundaries. Voxel boundaries and centres are in black, labelled *v*_1_ to *v*_4_. Nodes are in red; *n*_1_ to *n*_6_ are cortical (corresponding to surface vertices), *n*_7_ to *n*_10_ are subcortical (corresponding to voxels). Individual voxels may contain multiple nodes, which is an explicit representation of PVE.

An important property of hybrid space is that it provides an explicit representation of PVE. This follows from the fact that nodes in hybrid space are defined to only ever map to a single tissue type (which may be GM or WM, depending on where the node itself is). Voxels that are affected by PVE will therefore contain multiple nodes (for example, a subcortical WM node and some cortical GM nodes), as is the case for many of the voxels in figure 1. Toblerone can perform projection between the volume and surface spaces (a conventional mode of operation that is directly comparable with existing methods), or it can perform projection between the volume and hybrid spaces (a new capability for which there are no existing methods).

As nodes in hybrid space correspond to only one tissue, the signals associated with them must also respect this distinction. Signal on cortical nodes is considered as being purely cortical (i.e., GM) in origin, and signal on subcortical nodes is considered as purely subcortical in origin (i.e., WM or a subcortical GM structure). *How* these separate signals are derived from data that contains PVE is beyond the scope of this work (and is in fact the objective of partial volume effect correction, PVEc^3^). For the purposes of this work, it is only necessary to posit that the separate signals exist and that hybrid representation respects their separation. It follows that, when projecting from hybrid to volume space, the process of PVE must be recreated to mix the separate tissue signals within each voxel in the correct proportions.

### 2.2. Construction of the projection

The projection is constructed from mesh representations of the inner and outer surfaces of a cortical hemisphere, produced for example by FreeSurfer (in which case they are referred to as the white and pial surfaces respectively) [14]. Mesh representations consist of vertices (points in 3D space) and triangles (triplets of vertex numbers denoting how vertices should be connected to form the discrete primitives of the surface). Triangle correspondence between the inner and outer surfaces of the cortex is assumed. Though meshes of any resolution can be used, it is envisaged that meshes at 32k resolution are sufficient for physiological data (as is the case for HCP analysis pipelines [11]). Though the inner and outer surfaces of the cortex are used to define the extents of the cortical ribbon, the actual projection itself is made onto the cortical midsurface by default, though this could be modified for other use cases such as laminar fMRI. A set of PV estimates for the cortex is also used; these are produced using Toblerone’s previously-reported PV estimation function operating on the cortical surfaces [15]. Both the forward and reverse projections are constructed from the two matrices that are introduced in the following paragraphs.

#### Voxel-triangle matrix

The voxel-triangle matrix performs a mapping between individual voxels and the volume primitives that comprise the cortex. Construction of the primitives is approached in a triangle-wise manner, as opposed to the vertex-wise approach adopted by other methods, as this allows for the construction of extremely simple primitives. For each pair of triangles on the inner and outer surfaces, their corresponding vertices are connected to form a triangular prism^4^ which is subdivided into three non-overlapping tetrahedra [16]. For example, for triangles with vertices Δ*ABC* and Δ*abc* on the outer and inner surfaces respectively, the prism may be split into tetrahedra *ABCa, Cabc* and *BCab*. As a consequence of this subdivision, each quadrilateral face of the prism is split along the diagonal into two triangles. It is important to ensure that the orientation of this face diagonal is consistent between neighbouring prisms so as to avoid producing overlapping tetrahedra. This is achieved by fixing the direction of subdivision according to the ordering of vertex numbers: ‘positive’ if *A* < *B, B* < *C* or *C* < *A*, and ‘negative’ otherwise.

Subsequently, for each voxel in the neighbourhood of the prism, an isotropic grid of sample points is initialised. Figure 2 shows one such prism and the sample points from one neighbourhood voxel. Sample points are tested to determine if they lie within each tetrahedra (via the Delaunay triangulation [17]) and the results stored in the matrix VT. The elements VT_*v,t*_ are continuous values in the range [0,1] recording the fraction of sample points from voxel *v* that lie inside triangular prism *t*. This matrix is highly sparse: for a voxel size of 2.2mm isotropic and a surface resolution of 32k vertices, each triangular prism intersects around 10 voxels on average.

**Figure 2:**
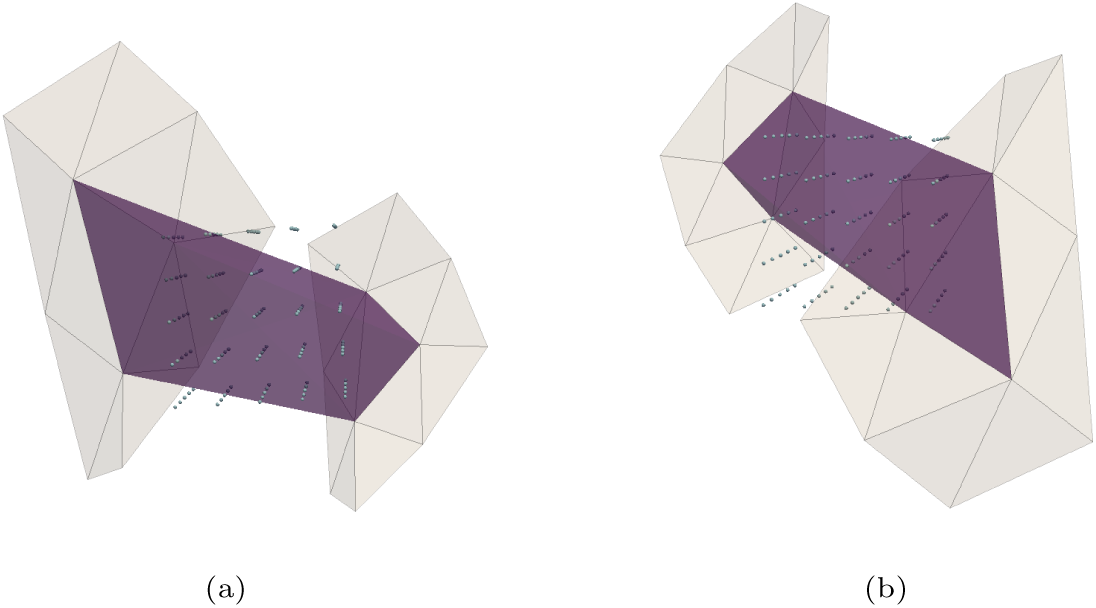
Two alternative views of the triangular prism (purple) formed by connecting one pair of corresponding triangles on the inner and outer surfaces (grey) of the cortex. The subdivision into three tetrahedra is not shown. A 5^3^ grid of sample points are tested to determine the volume of intersection between the voxel and prism, which is recorded in the voxel-triangle matrix.

#### Points-triangle matrix

The points-triangle matrix records the area of each triangle on the cortical midsurface that is associated to its individual vertices (though vertices are now referred to as points to avoid re-using the letter *v*). These values are calculated using the Voronoi region approach, extended to arbitrary meshes by Meyer *et al*. [18], and are recorded in the matrix PT. The element PT_*p,t*_ denotes the area of triangle *t* that is associated to point *p*. Once again, this matrix is highly sparse as each point is contained by only a few triangles.

#### Forward projection: volume to surface

To form the volume to surface projection, the VT matrix and PT matrices are first normalised in a row or column-wise sense, denoted with the ‖·‖_*row*_ operator for the row-sense. This operator rescales rows or columns with any non-zero elements such that the row or column sum is unity. For the VT matrix, this is equivalent to ensuring that the sum of voxel weights for each triangle is unity, and for the PT matrix, that the sum of triangle areas for each point is unity. The effect of this is to turn these individual matrices into weighted-averaging operators that preserve signal intensity. The forward projection is then defined by the multiplication

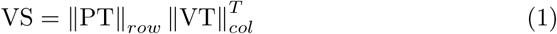

#### Forward projection

*volume to hybrid*. In order to extend the forward projection into hybrid space, it is necessary to include the mapping between voxels and subcortical nodes (the VS matrix accounts only for cortical nodes). This is achieved by concatenating the VS matrix with the identity I_*v*_ matrix of size (number of voxels) to yield VH. This implies a one-to-one mapping for all voxels to their hybrid space counterpart (it is assumed, but not required, that irrelevant voxels are masked out, in which case their corresponding nodes are also removed).

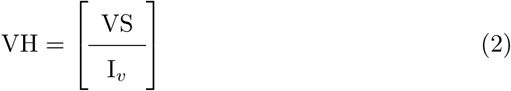

#### Forward projection: edge up-scaling

Depending on the nature of the data that is to be projected, edge up-scaling is an optional extra step that can be incorporated into the forward projection. For signals that scale directly with the amount of tissue in a voxel, signal magnitude will be artificially low in any edge voxels that are less than 100% brain tissue (for example, those that contain some CSF which does not contribute any signal). In this scenario, it may be advantageous to up-scale the signal to account for the lost intensity. This is performed by calculating the voxel-wise brain PV (the sum of GM and WM PVs), taking the inverse of this quantity to form an upweight vector **u**, and finally taking the outer product (denoted ⊗) with the constituent parts of the volume to hybrid projection matrix.

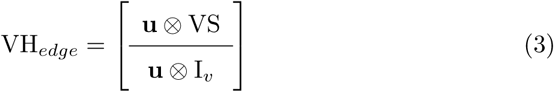

The purpose of this step is to perform a very simple correction for signal lost due to PVE, with the important caveat that it will be poorly-conditioned in voxels that contain a low GM and WM PV. Hence, this correction may not be appropriate for noisy data, but may be useful for data that has already undergone some form of processing. An example of where this step could be applied is projection of ASL data (the magnetisation signal scales with tissue PV); an example of where it would not be appropriate is projection of a timing volume (such as voxel-wise MRI slice times, which are independent of tissue PV).

#### Reverse projection: surface to volume

To form the surface to volume projection, the VT and PT matrices are normalised in the opposite sense to before. For VT, this ensures that the sum of triangle weights for each voxel is unity, and for PT, that the sum of vertex areas for each triangle is unity. The surface to volume projection is then defined by the following multiplication.

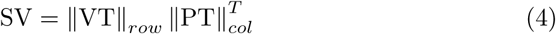

#### Reverse projection: hybrid to volume

In order to form the reverse projection from hybrid space, PVE must be recreated. This follows from the fact that nodes in hybrid space enforce the separation of signals from different tissues. Each row of the hybrid to volume projection matrix corresponds to a voxel, and the column values record the weights in which both cortical and subcortical nodes contribute to that voxel. A set of voxel-wise GM and WM PV estimates are then used to scale the overall contribution of cortical and subcortical nodes within each voxel, where **b** = **pv**_*gm*_ + **pv**_*wm*_ represents the voxelwise sum of PVs^5^. In particular, each row of the surface-to-volume matrix is scaled by the GM PV of the corresponding voxel, and each row of the identity by the WM PV of the corresponding voxel.

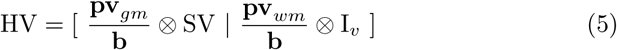

#### Reverse projection: edge down-scaling

Edge down-scaling may optionally be applied for the reverse projection depending on the nature of the data in question. The same principles as for the forward case apply: if the signal is expected to scale with tissue PV, then edge scaling reduces the projected signal in voxels that are less than 100% brain tissue. In this situation, there is a subtle distinction between PVE caused by *mixed* signal and PVE caused by *missing* signal, though both originate in the same manner (the presence, or absence, of multiple tissues in a voxel). Edge down-scaling is achieved by assembling the constituent parts of the hybrid to volume projection matrix without the weighted averaging implied by the brain PV **b**.

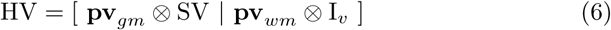

Though existing methods have largely focussed on the forward projection only (from volume to surface), the reverse projection does have some useful applications. For example, it can be used to generate *in silico* phantoms: starting from separate sets of parameter values for the cortex and subcortex, the reverse projection can be used to generate the corresponding volumetric data that one would expect to acquire for a given anatomy, incorporating the effect of PVE.

## 3. Materials and methods

### 3.1. Comparator methods

Toblerone’s projection was compared against the HCP’s ribbon-constrained (RC) method, which was developed for use with BOLD data in the HCP’s *fMRISurface* pipeline [11]. The RC method assumes vertex correspondence between the inner and outer surfaces of the cortex. For each vertex in turn, the outermost edges of the triangles that surround that vertex are connected between the two surfaces to form a volume primitive (polyhedron) enclosing a small region of cortex. Nearby voxels are subdivided isotropically and the subvoxel centres tested to determine if they lie interior to the polyhedron. The interior fraction gives a set of kernel weights that are used to perform weighted-average mappings from voxels to each surface vertex in turn. The reverse projection may readily be constructed from the same set of kernel weights.

Implementations of the methods introduced by Grova *et al*. and Operto *et al*. [9, 10] were not available and so these were not investigated (nor do these methods explicitly define a reverse projection). In the spirit of the inverse matrix approach introduced by Lonjaret *et al*. [19], two extra variants of Toblerone and RC were investigated, entitled Toblerone-lsQR and RC-lsQR. For any projection of the form **y** = A**x** (be it surface-volume or volume-surface), the inverse projection can be found by solving the equation for **x** using the lsQR algorithm, which is able to solve sparse systems that are both under-or over-determined [20]. For experiments involving a round-trip of projection (introduced in section 3.2.1), the lsQR variants of each method were run at the intermediate step as a means of completing the round-trip. For example, when using Toblerone to perform a surface (A) to volume (B) to surface (C) projection, the Toblerone-lsQR variant was constructed by using lsQR on the intermediate data B in order to reach C. It is important to note that lsQR does not yield an inverse projection in the general sense; rather, for a specific vector of projected data, it is able to invert the projection *for that data only*. This is contrast to Toblerone and RC that define all of their projections independently of the data to be projected.

Comparisons were performed in both the forward (volume to surface) and reverse (surface to volume) directions. Where possible, differences between the methods were tested for statistical significance using a paired *t*-test of independent measures where equal variances were not assumed and the significance threshold was set at *p* = 0.01.

### 3.2. Datasets

Two types of data were simulated, with local and global signal distributions respectively. For both data, the underlying anatomical information was taken from the structural pre-processed output of the 45 subject test-retest cohort of the Human Connectome Project, onto which physiological data were simulated [11]. Left white and pial cortical surfaces at 32k mesh resolution were used to define cortical anatomy. Whilst Toblerone is able to perform simultaneous projection to the left and right cortical hemispheres in a single operation, the RC method is not, and hence the comparison was restricted to consider single hemispheres only.

#### Local cortical signal

The first dataset, defined for the surface space only, models signals that are local in nature. The principle, previously used by both Grova and Operto, was to simulate small areas of activation within the cortex by selecting a random vertex on the cortical midsurface (with local thickness greater than 2 mm to avoid areas with high partial voluming), and applying the following signal values to the point, its first-order neighbours, second-order, and so on: [1, 1, 0.9, 0.7, 0.5, 0.3, 0.1]. All other vertex values were left at zero. Unlike the work of Grova and Operto, the location of the activation was allowed to vary between subjects; one such activation is shown in figure 3. Toblerone (in surface/volume mode) and RC were used on this dataset.

**Figure 3:**
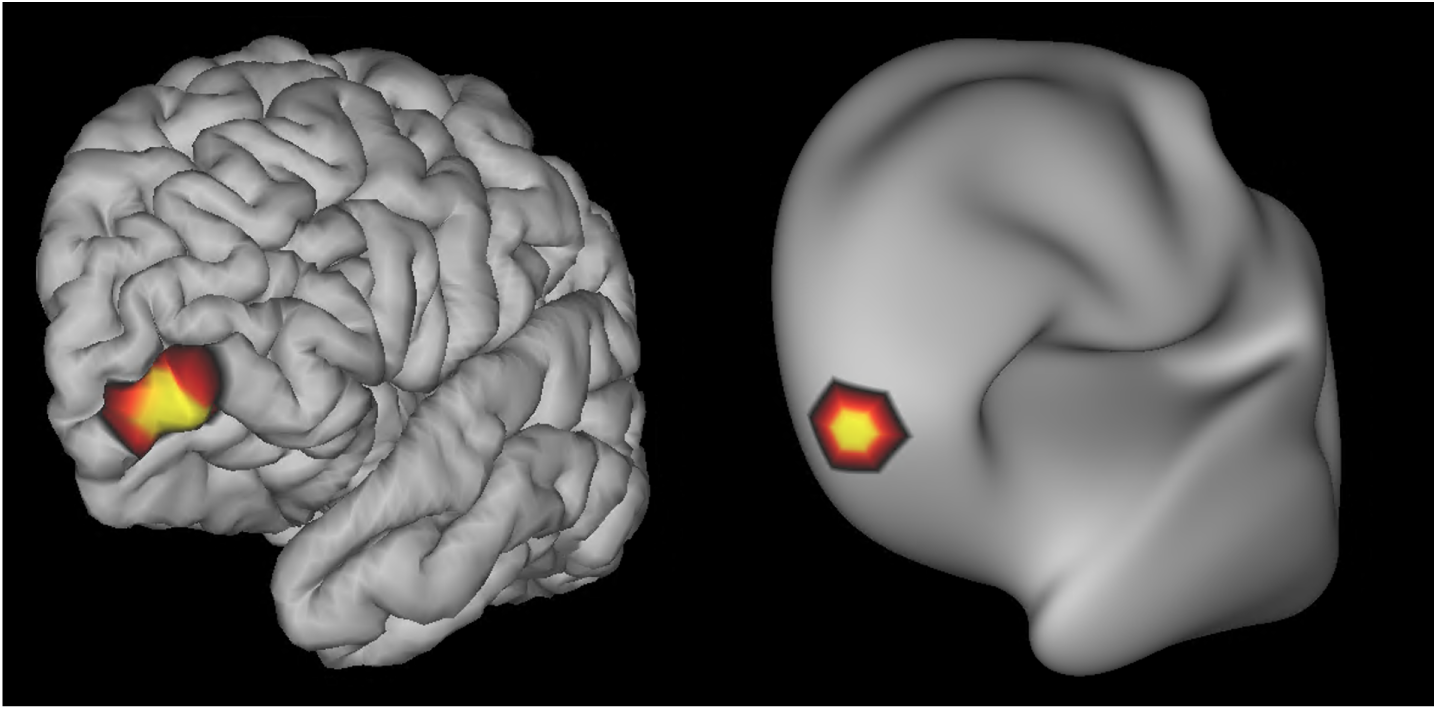
Example activation for subject 103818 of the HCP. The same signal map is shown on both the pial surface (left) and inflated surface (right).

#### Global cortical signal

The second dataset, defined for both the surface and volume spaces, models a global signal in the cortex only. Two different variants were tested: one with a uniform signal of 60 a.u., and one with sinusoidal variation of 60 ± 18 a.u. The volumetric data were produced by evaluating volumetric 3D fields of values and then performing a voxel-wise multiplication with subject-specific cortical GM PV estimates obtained via Toblerone’s surface-based PV estimation. In order to produce surface data, trilinear interpolation of the same volumetric fields onto each subject’s midsurface was performed. This process is illustrated in figure 4. Toblerone (in surface/volume mode), RC, Toblerone-lsQR and RC-lsQR were used on this dataset.

**Figure 4:**
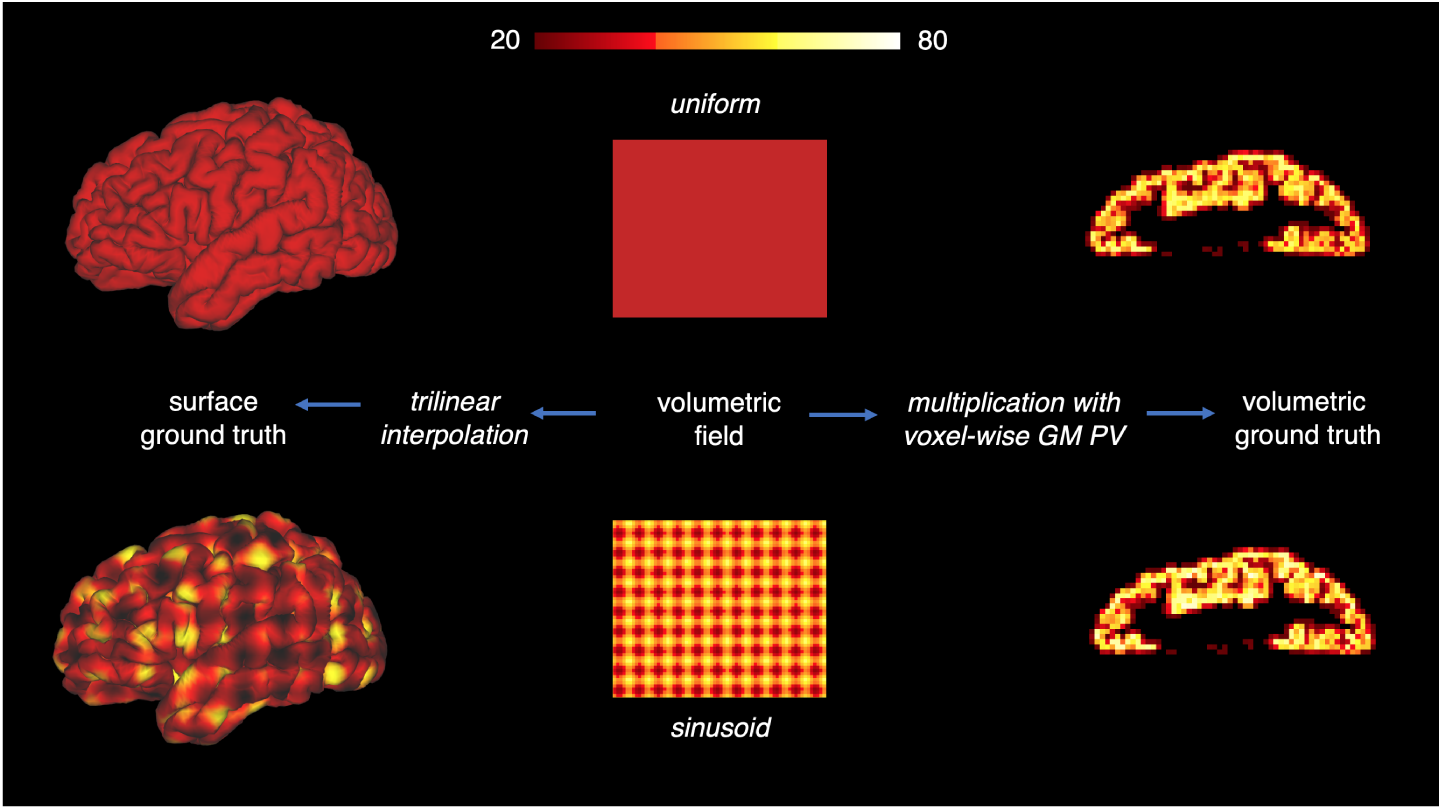
Generation of ground truth surface and volume data for both uniform and sinusoid signals.

#### Global cortical and subcortical signal

The final dataset, defined for both the surface and volume spaces, models a global cortical and subcortical signals. The cortical signal was generated in the same manner as for the global cortical signal dataset, onto which the following subcortical signals were added. For the uniform dataset, a constant value of 20 in all of the subcortex, and for the sinusoidal variation, a field of 20 ± 2. In the volume space, the cortical and subcortical signals were combined in proportion to voxel-wise GM and WM PV estimates. In the hybrid space, the ground truth was generated by concatenating a vector of subcortical signal values to the vector of cortical values. For simplicity, the subcortex was taken to contain only WM and no deep subcortical GM structures; in any case, because the projections consider only voxels intersecting the cortex, this anatomical inaccuracy is of no consequence. Toblerone (in hybrid/volume mode), RC, Toblerone-lsQR and RC-lsQR were used on this dataset, though the RC methods deal only with surface nodes and are unaware of the presence of subcortical nodes, so the comparison is incomplete.

##### 3.2.1. Experiments

Due to a lack of ground-truth projection methods, it is difficult to experimentally determine if one projection is more correct than another. Instead, other metrics of performance must be evaluated, for example area under receiver operator characteristic curve (AUROC) of activation detection which has previously been used for this application. A novel aspect of this work is the extensive use of round-trip projections: projection of data to one space, followed by inverse projection back to the first space and comparison with the original data to check for similarity. This is possible because all the methods investigated define a reverse projection. The assumption behind this experimental design is that a projection that is more self-consistent on a round-trip is also likely to be more performant in other ways.

#### Local cortical signal

The ground truth surface activations were taken through a round-trip of projection, from surface to volume and back to surface, and the result run through a binary classifier to identify activated and unactivated vertices. AUROC analysis, as used by Grova and Operto [9, 10], was used in this work, which includes a normalisation step such that the range of the projected data is the unit interval. For the dataset in question, the number of true negative vertices vastly exceeds the number of true positives, which biases the analysis, and hence the AUROC_close_ variant introduced by Grova was used here [9]. For each activated vertex on the ground truth, a corresponding unactivated vertex within the fourth-order neighbourhood of vertices was selected, and the classification performed on the evenly weighted sample of vertices. Restricting the sampling to a close neighbourhood ensures falsely activated vertices are included which renders the classification more challenging.

AUROC analysis, as performed above, operates on a relative basis due to the normalisation step. Whilst this is not unreasonable for the analysis of BOLD data, with ASL it is typically desired to perform absolute quantification and hence normalisation is inappropriate. A parallel analysis of the same dataset was thus performed on round-trip projected data without normalisation and three metrics evaluated. Firstly, the maximum signal value in the volume space, which is indicative of the PVE-induced scaling incorporated into each projection method. Secondly, the maximum signal value when projected back into the surface space. Though neither method will be able to perfectly reproduce the ground truth (due to PVE), the extent to which signal intensity is preserved is nevertheless important for absolute quantification strategies^6^. Finally, the ratio of activated vertices to ground truth activated vertices after the roundtrip projection, as this indicates the extent to which the projection has smoothed the original signal distribution.

#### Global cortical (and subcortical) signal

These experiments were run on the global cortical signal data, with and without subcortical signal. Surface and volumetric ground truths were taken through a round-trip of projection, as illustrated in figure 5. The lsQR variants of each projection were also used at the intermediate step to obtain the inverse projection. The following metrics were calculated with respect to ground truth: root-mean-square (RMS) error, to give an overall indication of how accurately ground truth was reproduced; and minimum and maximum signal values, to investigate how signal intensity and range was preserved. The latter metrics are of interest because an ideal projection should not produce signal values that are outside of the input signal range. In the context of these experiments, where the ground truth data were always positive, this means projection should never yield a negative signal value, or value above 100% of the ground truth maximum.

**Figure 5:**
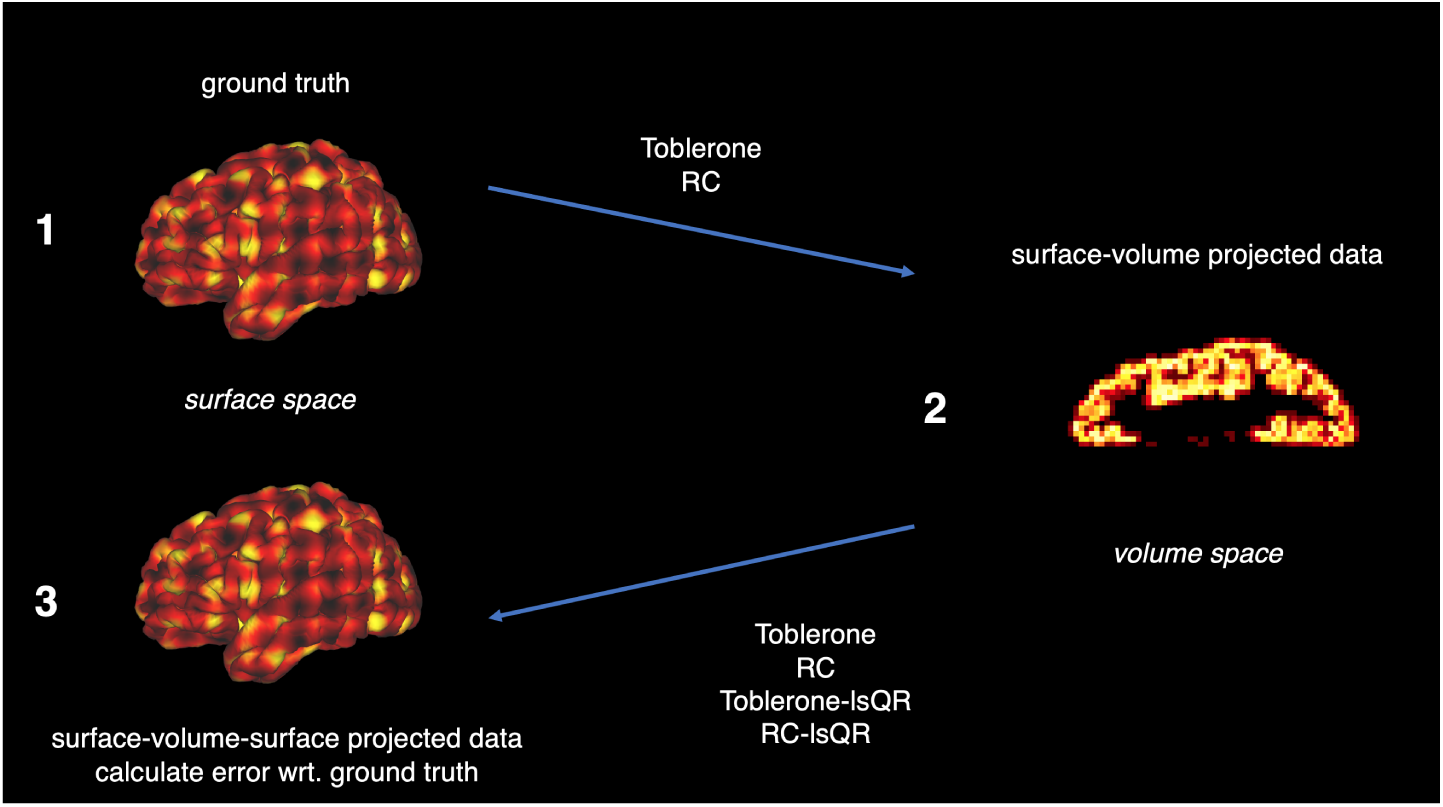
Schematic of the surface to volume to surface round-trip projection experiment. Both Toblerone and RC were used to project ground truth surface data into the volume. The volumetric data were then projected back to the surface using two methods: each method’s self-inverse (*i.e*., Toblerone’s volume to surface), or the lsQR inversion of each method’s forward projection (*i.e*., Toblerone-lsQR in this case). Finally, metrics were calculated with respect to ground truth in the volume space. The volume to surface to volume experiment was of the same form, with the spaces swapped around.

### 3.3. Ethics statement

As this work is computational methods development using previously published public datasets, no further ethical review was performed besides that of the original Human Connectome Project [12].

## 5. Results

### 5.1. Local cortical signal

Figure 6 shows the AUROC analysis on round-trip projected local cortical signal surface data with normalisation. Both methods returned extremely similar outputs, to the point where no statistically significant difference could be detected.

**Figure 6:**
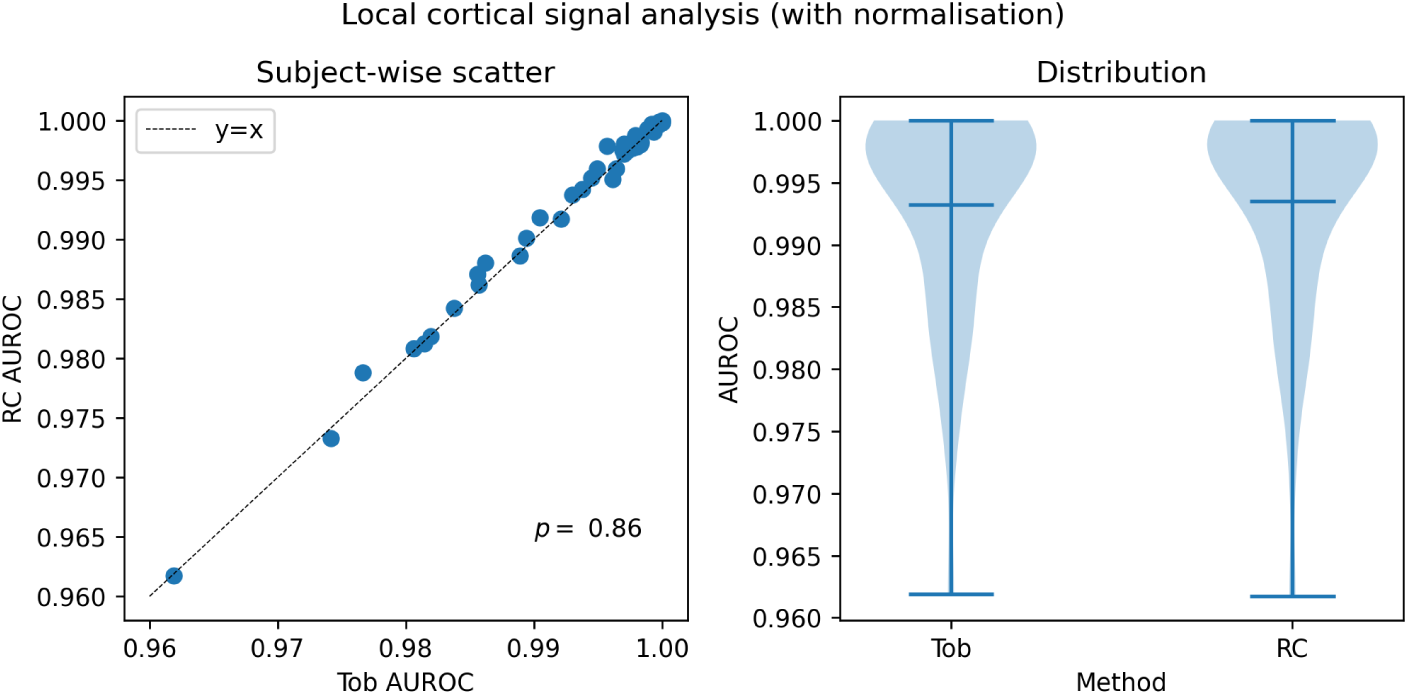
AUROC scores for the round-trip projected local cortical signal data. Right: a per-subject scatter plot of scores revealed no statistically significant difference between the two methods. Left: both methods returned extremely similar distribution of scores across subjects.

Figure 7 shows the analysis of round-trip projected local cortical signal data without normalisation. Both methods returned extremely similar distributions of maximum signal values for projection into the volume or back onto the surface, though in both cases there was a small and statistically significant advantage in favour of Toblerone. When considering the number of non-zero vertices following round-trip projection, expressed as a fraction of the ground truth, Toblerone returned a distribution slightly wider and with higher mean than RC.

**Figure 7:**
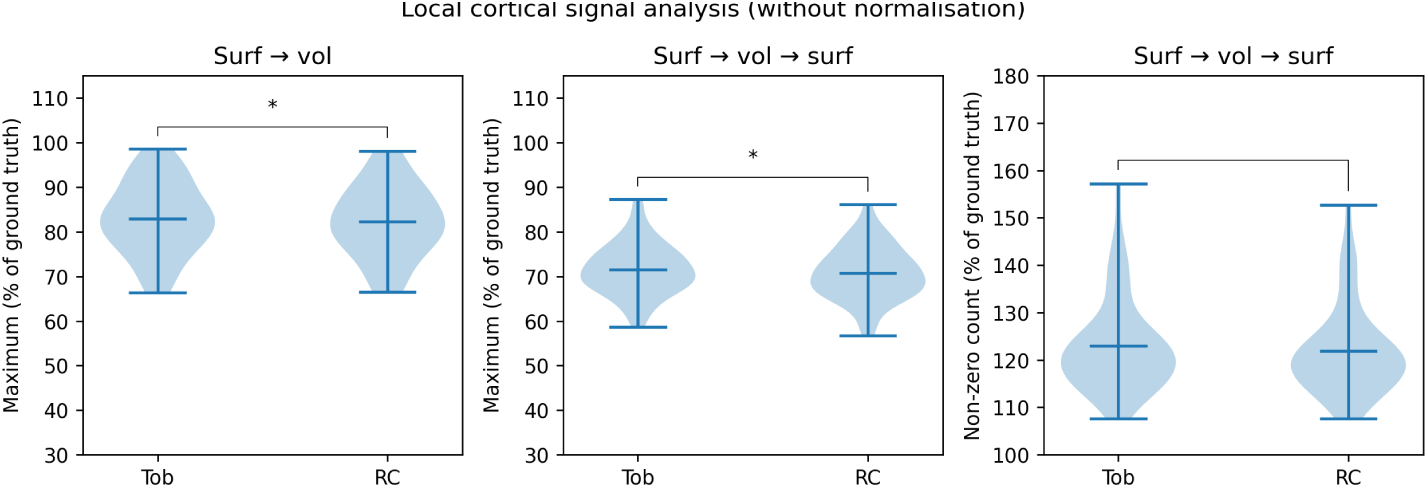
Round-trip projected local cortical signal surface data without normalisation. Left: distribution across subjects of maximum signal value after projection into volume space. Centre: distribution across subjects of maximum signal value after projection back onto the surface. Right: the distribution across subjects of non-zero vertices, as a proportion of ground truth, was slightly wider and of greater mean for Toblerone, though the difference was not statistically significant. *\** denotes a comparison of statistical significance.

### 5.2. Global cortical signal

Figure 8 shows the results of the round-trip projection on global cortical signal data, starting and ending in the volume space. In terms of RMS error, Toblerone performed better than RC (statistical significance was reached), whereas for maximum signal values both methods performed similarly. By contrast, lsQR methods returned essentially ‘ideal’ values for RMS error and maximum signal, though at the cost of slightly negative minimum signal values.

**Figure 8:**
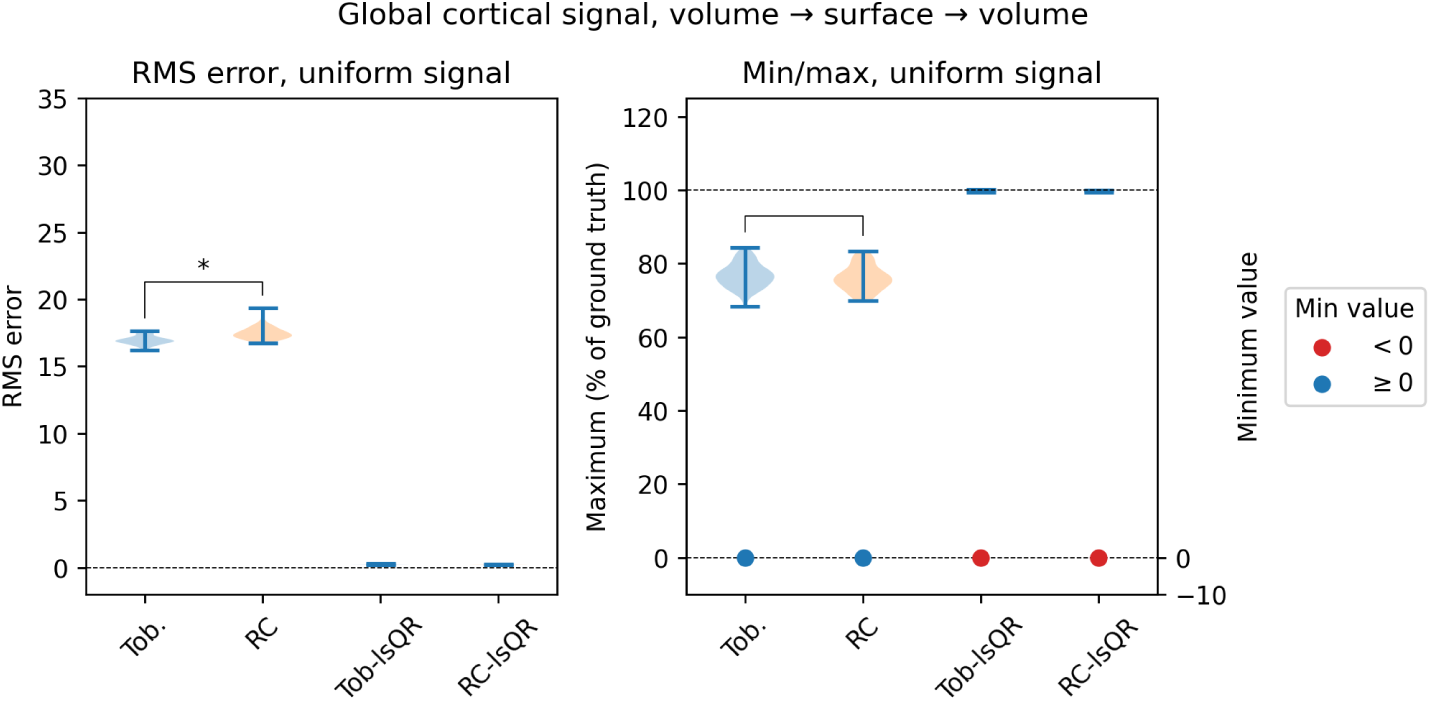
Global cortical signal, uniform value, round-trip projection from volume to surface to volume. Left: Toblerone produced significantly lower RMS errors than RC of around 17, whereas lsQR methods returned almost no error. Right: both Toblerone and RC returned similar maximum signal values of around 80%, with no significant difference between the two, and both returned zero signal minimums. lsQR methods returned almost maximum signal values of almost 100% of ground truth, but the minimum signal values were slightly negative. *\** denotes a comparison of statistical significance.

The complete results from this experiment (given in the supplementary material) were in agreement with the subset presented in figure 8: the RMS errors produced by Toblerone were always somewhat lower than RC (statistical significance was reached); and the maximum values produced by both methods were essentially identical (no significant difference observed). lsQR methods produced lower RMS errors than their corresponding original method (ie, ToblsQR lower than Toblerone itself); but minimum and maximum signal values for these methods were observed to fall outside the desired range of zero to 100% of ground truth.

### 5.3. Global cortical and subcortical signal

Figure 9 shows the results of the round-trip projection on global cortical and subcortical sinusoid variation data, starting and ending in the surface space. Toblerone produced lower RMS errors than RC (statistical significance was reached); both lsQR methods returned lower errors but were still substantially greater than zero. Both Toblerone and RC returned maximum signal values around 80% of ground truth, whereas lsQR methods returned a wide range of values that were often greater than 100%. All methods returned minimum values somewhat above zero.

**Figure 9:**
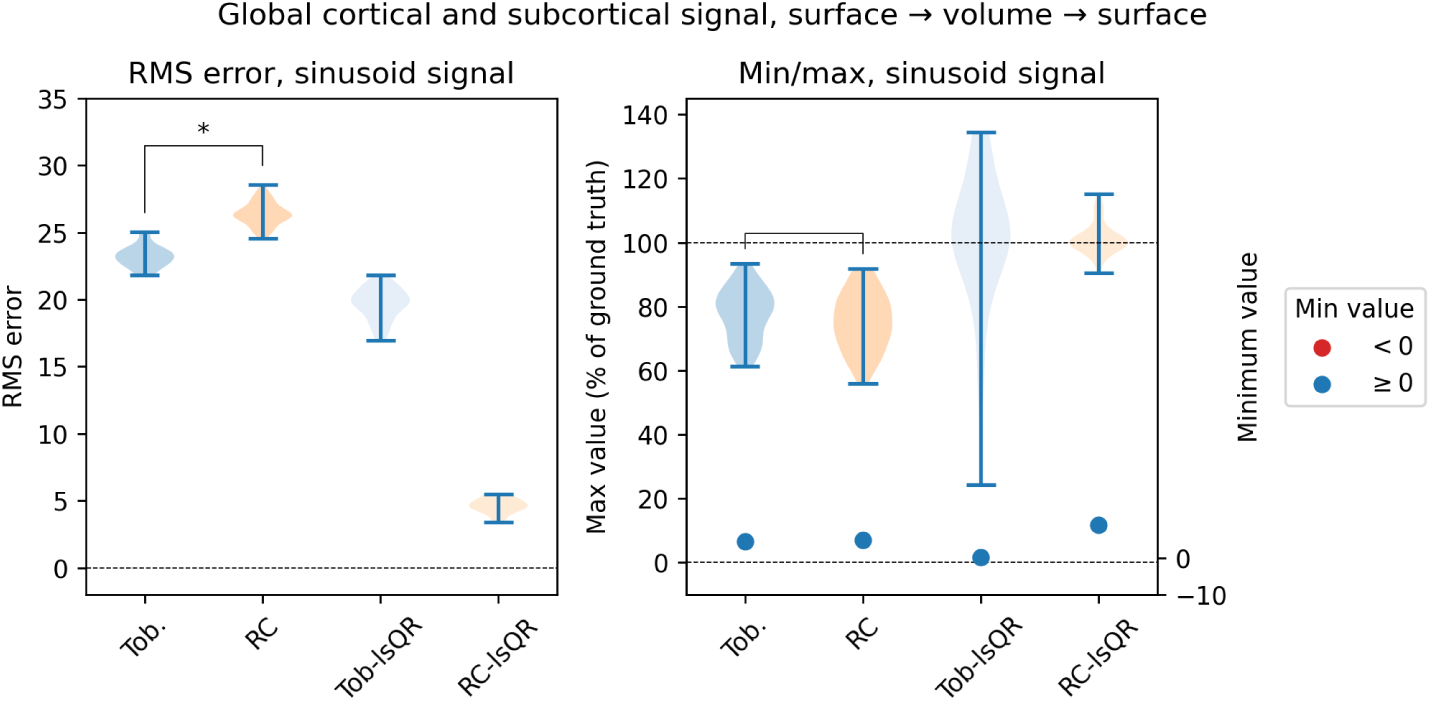
Global cortical and subcortical signal, sinusoid variation, round-trip projection from surface to volume to surface. Left: Toblerone returned significantly lower RMS errors than RC (around 24 versus 26 a.u.). RC-lsQR performed substantially better than Tob-lsQR, and both lsQR methods performed better than their original counterparts. Right: maximum and minimum signal values. Toblerone and RC returned similar maximums, with minimums somewhat above zero. lsQR methods, particularly Tob-lsQR, returned maximum signal values substantially above 100% of ground truth, up to 130% in the extreme case. *\** denotes a comparison of statistical significance.

**Figure 10:**
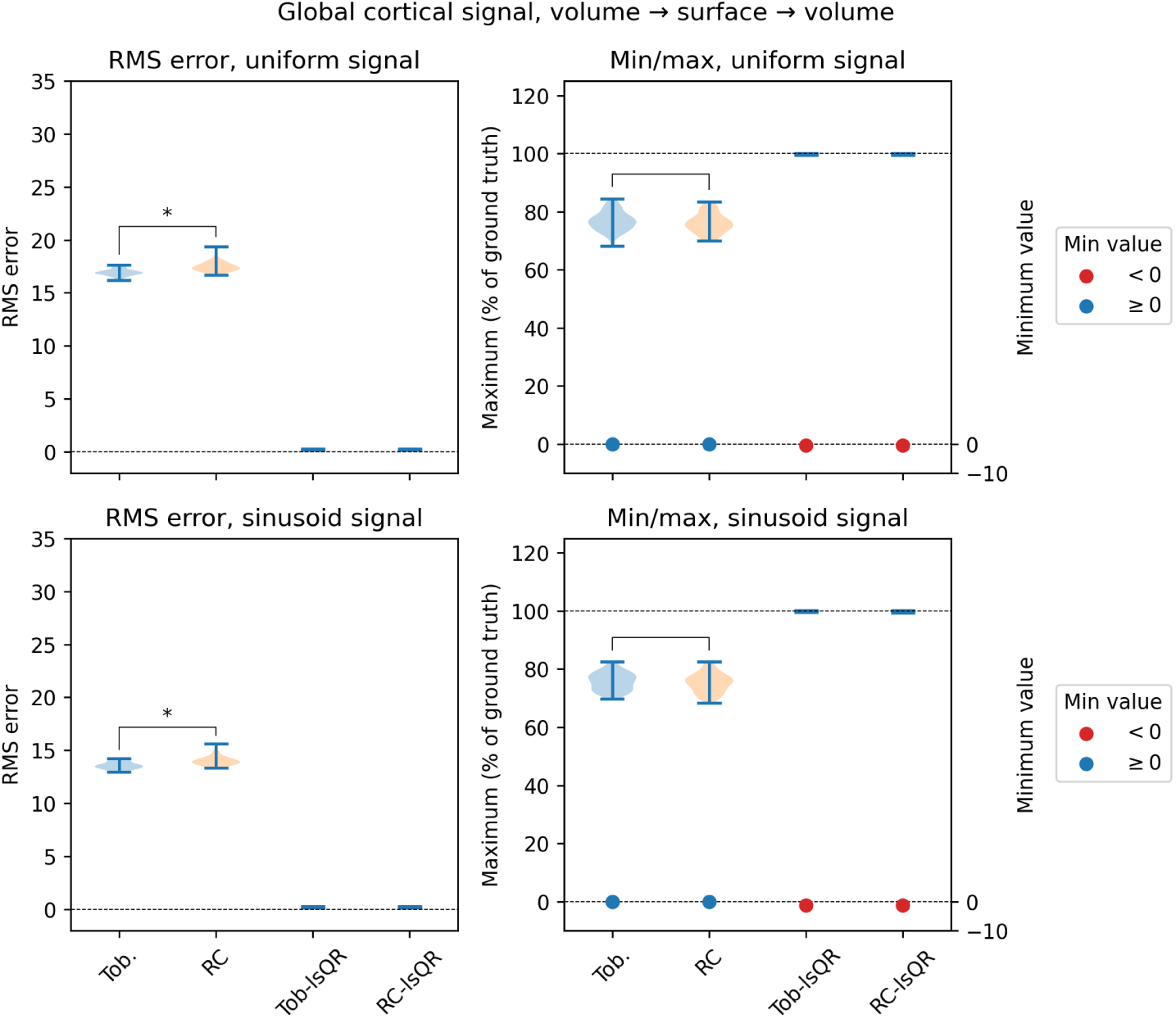
*\** denotes a comparison of statistical significance.

The complete results from this experiment (given in the supplementary material) were in agreement with the subset presented in figure 9: the RMS errors produced by Toblerone were always notably lower than RC (statistical significance was reached); and the maximum values produced by both methods were essentially identical (no significant difference observed). lsQR methods produced lower RMS errors that their corresponding original method (ie, Tob-lsQR lower than Toblerone itself); but minimum and maximum signal values for these methods were observed to fall outside the desired range of zero to 100% of ground truth.

## 6. Discussion

The results of the local cortical signal tests, both using AUROC analysis and without normalisation, showed that Toblerone produced extremely similar outputs to RC on a conventional evaluation of projection. Across all analyses, the most notable difference between the two was the slightly higher smoothing of signal produced by Toblerone (right panel of figure 7), though this was not statistically significant. It was therefore concluded that the methods produced similarly for conventional volume and surface projection, which is expected given the closely related geometric foundations upon which they are built.

The results of global cortical signal tests, both with and without a subcortical signal (figures 8 and 9 respectively), demonstrated more substantial differences in behaviour between Toblerone and RC. On all four variants of this experiment, Toblerone produced significantly lower RMS errors than RC following a round-trip of projection, with the differences largest on those experiments involving data with a subcortical signal. Whilst neither method could ever produce zero error (because the down-scaling caused by PVE in the surface to volume direction is not reversed in the volume to surface direction) Toblerone’s lower errors show that it is more self-consistent than the RC method. Regarding minimum and maximum signal values, both Toblerone and RC returned very similar results, which was in line with their results on the local cortical signal tests. Importantly, though neither method was able to perfectly recover ground truth signal values, they always mapped them to a desired range of zero to 100% of ground truth: no negative values were produced, nor values above the original maximum. It is acknowledged that the comparison with RC on data containing subcortical signal is imperfect, because RC is not designed for this purpose. Nevertheless, the results of the comparison demonstrate why existing projection techniques are unsuitable for data that contains a subcortical signal of interest.

The lsQR methods performed extremely well on some isolated experiments, notably those of the volume to surface to volume design, for which lsQR was used to perform the latter surface to volume projection by inverting the prior volume to surface output. It is likely that good performance for this step was due to the surface to volume projection being over-determined (there are multiple surface vertices per voxel) and hence the matrix inversion is favourable. By contrast, when lsQR was used to perform a volume to surface projection, which is under-determined, higher RMS error and numerical instability was observed (see figures 11 and 13). In particular, lsQR methods returned maximum and minimum signal values outside of the desired range of zero to 100%: both negative values and values over 100% of ground truth were observed. These findings show both the promise and pitfalls of working with an inverse-based method: the potential for excellent recovery of ground truth (including the removal of PVE scaling effects), but at the risk of numerical instability leading to signal values far outside of the desired range, which would be a substantial hinderance for any quantitative analysis strategy. It is for this reason that Lonjaret *et al*. devoted significant effort in their work to regularising the inversion step, but in doing so made the method specific to BOLD [19].

**Figure 11:**
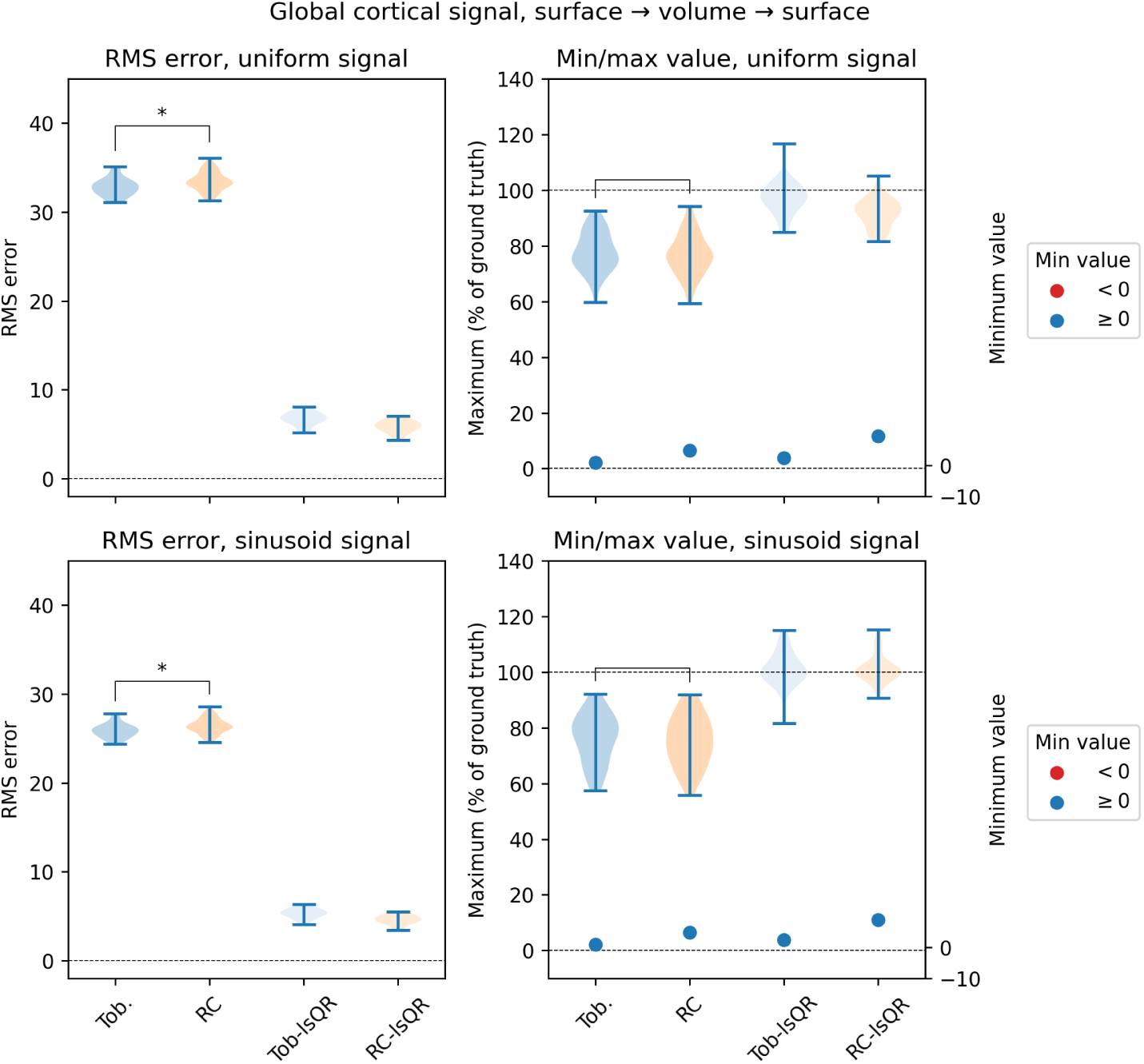
*\** denotes a comparison of statistical significance.

**Figure 12:**
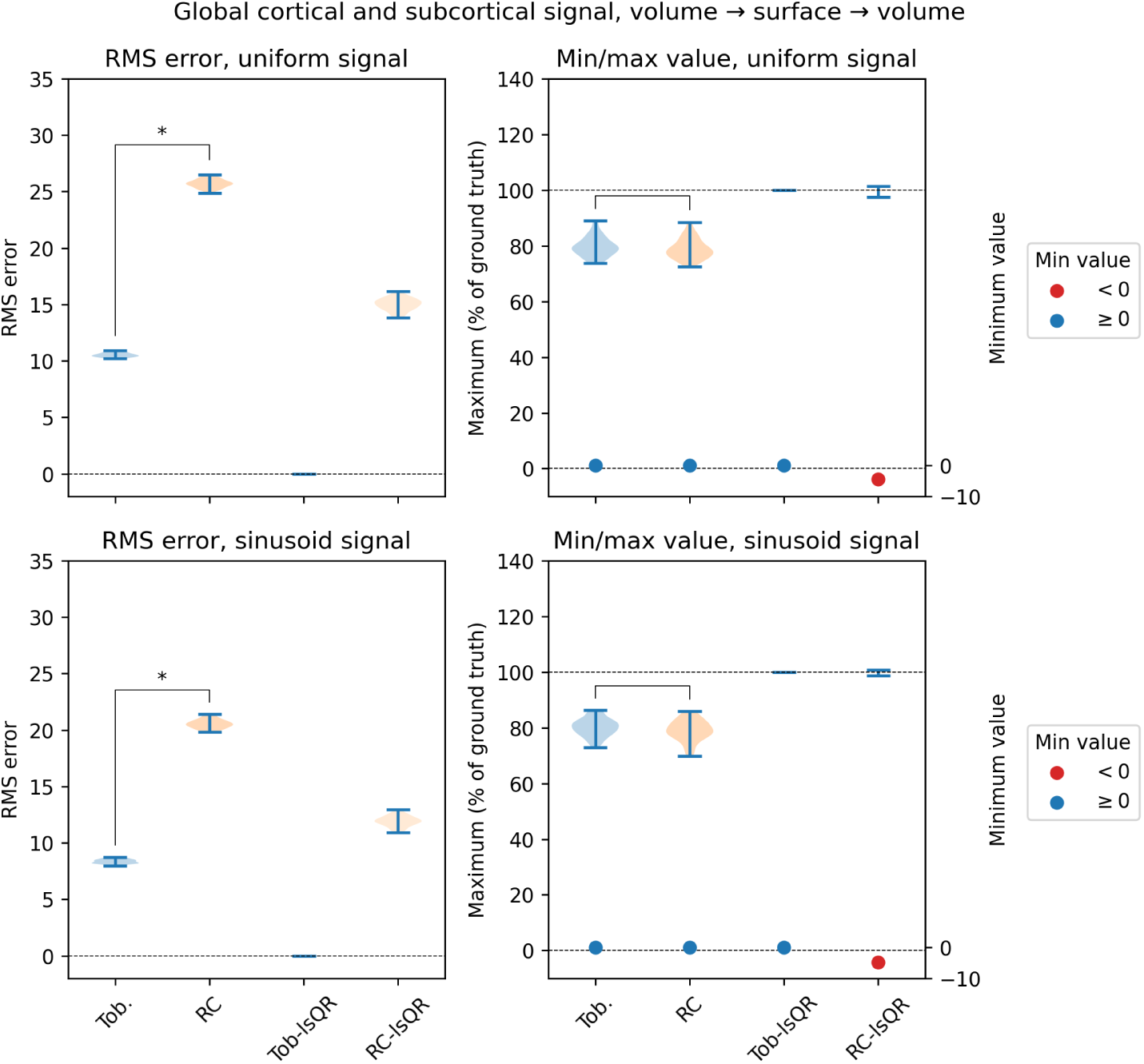
*\** denotes a comparison of statistical significance.

**Figure 13:**
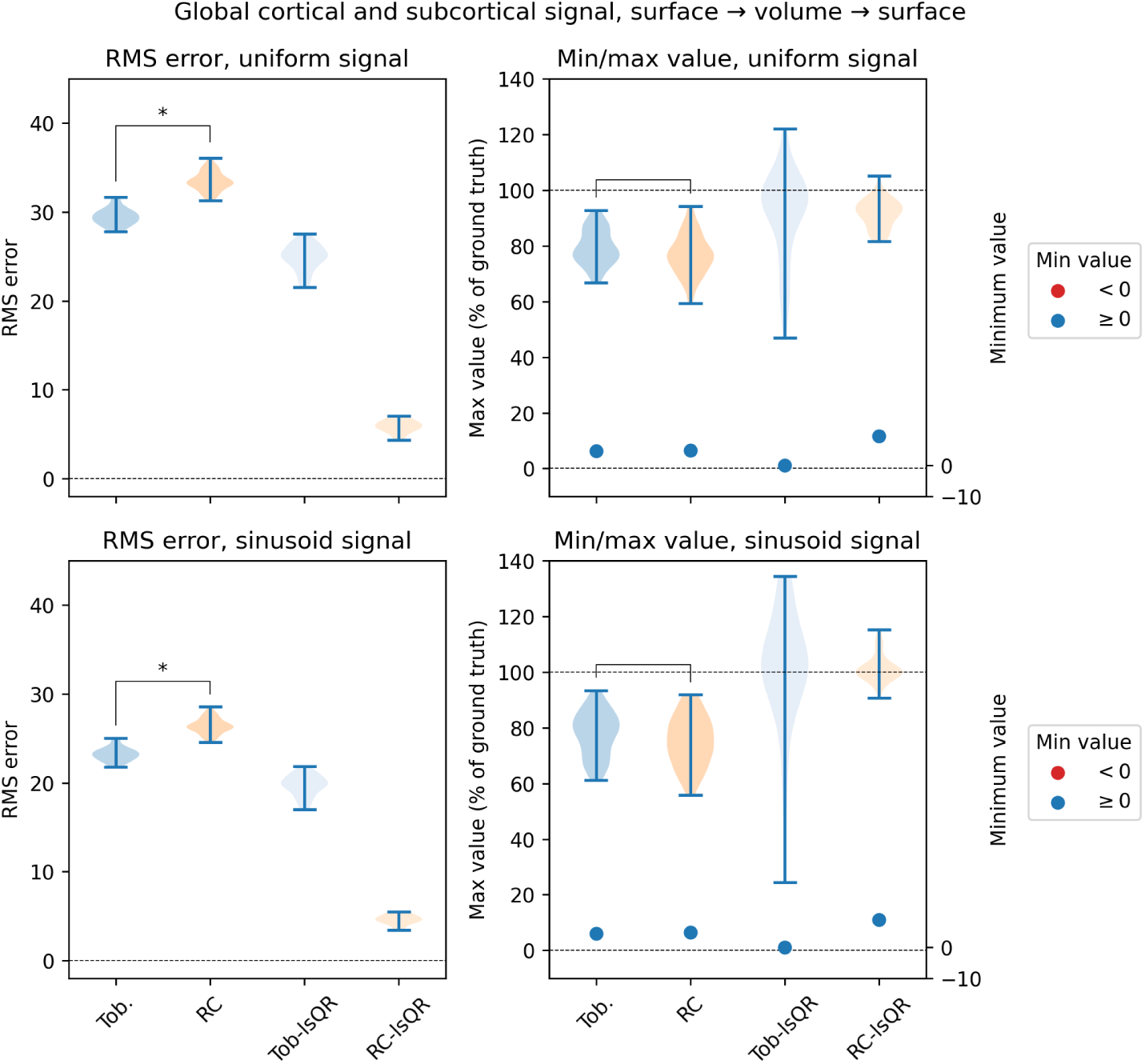
*\** denotes a comparison of statistical significance.

## 7. Conclusions

Toblerone is an extension and generalisation of existing approaches to projection that is able to work with arbitrary data, without making modality-specific assumptions, and project to and from hybrid space, a close analogue of the HCP’s grayordinates space. The geometric construction of the projection is closely related to the HCP’s RC method, and accordingly we observed that the two methods produced similar outputs when operating in a conventional volume-surface sense on data that contains only cortical signal. When operating in hybrid space on data that contains both cortical and subcortical signal, larger and statistically significant differences compared to RC were observed. This implies that the novel aspects of Toblerone’s projection are of consequence and make it better suited to working with such data.

## 8. Acknowledgements

The authors would like to thank Tim Coalson of Washington University in St Louis, USA, for advice concerning the correct ordering of triangle vertices. Funding was provided by the EPSRC (EP/P012361/1) and (for T. F. Kirk) the Bellhouse scholarship at Magdalen College, Oxford. HCP data was provided by the Human Connectome Project, WU-Minn Consortium (Principal Investigators: David Van Essen and Kamil Ugurbil; 1U54MH091657) funded by the 16 NIH Institutes and Centers that support the NIH Blueprint for Neuroscience Research; and by the McDonnell Center for Systems Neuroscience at Washington University.

## 10. Data availability

The Toblerone software package is publicly available at https://github.com/tomfrankkirk/toblerone. The analysis code used to perform the experiments presented in this work may be accessed at https://github.com/tomfrankkirk/projection_paper. Finally, the structural data on which these experiments were performed are available directly from the HCP at https://db.humanconnectome.org.

## 11. Supplementary material

## Footnotes

1 The CIFTI file format embodies this dual representation explicitly [13].

2 https://github.com/tomfrankkirk/toblerone

3 Though many projection methods acknowledge the existence of PVE within data, there is little that can be done to remove this during the narrow operation of projection. To do so would require a basis on which to separate cortical and subcortical signals, which is a modality-specific consideration and explains why PVEc is a non-trivial operation.

4 These are similar to a Toblerone bar, hence the name.

5 Note the double-scaling here, which reflects the slightly different types PVE at work. Firstly, there is the question of determining the relative contributions of the different cortical nodes within the voxel (a weighting between neighbouring GM nodes); secondly, there is the question of determining the relative contributions of the cortical versus subcortical signals (the overall ratio of GM to WM).

6 Though Toblerone does implement edge up-scaling, this was not included in the interests of a like-for-like comparison with RC.

